# Intraperitoneal injection of sodium pentobarbital is associated with pain in rats

**DOI:** 10.1101/703173

**Authors:** JN Reimer, C Schuster, CG Knight, DSJ Pang, VSY Leung

**Affiliations:** Veterinary Clinical and Diagnostic Sciences, Faculty of Veterinary Medicine, University of Calgary, Calgary, AB, Canada; Department of Clinical Sciences, Faculty of Veterinary Medicine, Université de Montréal, Saint-Hyacinthe, QC, Canada; Groupe de Recherche de Pharmacologie Animale du Québec (GREPAQ), Faculty of Veterinary Medicine, Université de Montréal, Saint-Hyacinthe, QC, Canada

**Keywords:** euthanasia, nociception, writhing, grimace scale, behavior

## Abstract

An effective and pain-free killing method is required to achieve the goal of euthanasia, a “good death”. Overdose of sodium pentobarbital (PB) by intraperitoneal (IP) injection is a widely accepted technique, but questions remain regarding pain associated with administration. As PB rapidly causes sedation and loss of consciousness, most studies have relied on indirect evidence of pain. The objective of this study was to assess pain associated with IP PB using an appropriate vehicle control.

Adult male and female Sprague Dawley (SD) and female Wistar rats (N = 112) were block randomised by sex and strain to receive one of four treatments: 1) 800 mg/kg PB (pH 11); 2) 800 mg/kg PB with 4 mg/kg lidocaine (PB+lido); 3) saline or 4) vehicle controls (pH 11 or 12.5). Behavior (Rat Grimace Scale [RGS], writhing, back arching) was evaluated at baseline, before loss of righting reflex (PB and PB+lido groups), 80s, 151s and 10 min post-injection (PI; saline and vehicle control groups).

In the vehicle control groups, the RGS scores were increased at 151s PI (SD: p = 0.0008, 95%CI −0.731 to −0.202) from baseline, as was relative frequency of writhing (SD: p < 0.00001; Wistar; p = 0.0004). RGS scores remained elevated 10 mins PI (SD: p = 0.0070, 95%CI −0.768 to −0.118; Wistar: p = 0.0236, 95%CI −0.907 to −0.0742) but the relative frequency of writhing did not (p > 0.05). The RGS scores and the relative frequency of writhing remained low in the PB, PB+lido and saline groups (p > 0.05). Back arching increased from baseline in the PB+lido group before loss of righting reflex and in the vehicle control group (SD rats) at 151s PI (p < 0.05).

These results show that IP PB results in signs associated with pain. The sedative effects of PB limit behavioral assessment.

## Introduction

Approximately 9 million mice and rats are used in biomedical research in Canada and the European Union annually. [1, 2] As the majority of animals are killed at project completion (or when a humane endpoint is reached), an effective, fast and pain-free killing method is essential.

Two methods are generally considered to be acceptable: 1) injection of barbiturates, such as sodium pentobarbital (PB), and 2) an overdose of an inhalant anesthetic. [1, 3] The use of inhalant anesthetics for euthanasia has been reported as aversive to rodents. [1, 4, 5] Therefore, the intraperitoneal (IP) injection of an overdose of PB is widely considered to be a preferable method of euthanasia. [1, 3] The effect of IP PB has been reported to be quick with loss of righting reflex (LORR) and cessation of heart beat (CHB) occurring within approximately 111s and 283s, respectively, with 800 mg/kg PB. [6] However, it has been suggested that the highly alkali pH of PB may cause pain upon injection [7] and current guidelines recommend the use of local anesthetics, such as lidocaine, in conjunction with the IP injection of PB to minimize this effect. [1, 3] Few studies have explored the potential for pain or defined methods to assess pain associated with this killing method to support the addition of local anesthetics to the injectate. Studies that have explored pain associated with IP injection of PB report that while some behaviors associated with pain increase (e.g. writhing) others remain unchanged (e.g. the Rat Grimace Scale (RGS)). [8–10] Importantly, these studies did not account for the sedative effects of PB and the potential to interfere with behaviors used to evaluate pain, which may explain these conflicting results. The aim of this study was to assess if the injection of a vehicle control (with a similar pH to PB) is painful. It was hypothesized that pain behaviors would increase after the injection of a vehicle control but not following injection of PB.

## Methods

### Ethical statement

This study was approved by The Health Sciences Animal Care Committee at the University of Calgary (Animal Use Protocol AC11−0044) and was performed in accordance with the Canadian Council on Animal Care Euthanasia Guidelines (2010) and the Canadian Association of Laboratory Medicine (CALAM) Standards of Veterinary Care (2007).

### Experimental Animals/Housing and Husbandry

Adult male (n = 48, 359g [201 to 440g] [median, range]) and female (n = 53, 263g [196 to 448g]) Sprague Dawley (SD) surplus rats were obtained from the University of Calgary Health Sciences Animal Resources Centre (HSARC). Female Wistar rats (n = 50, 239.5g [212 to 265g]) were obtained from Charles River Canada. Animals were housed in pairs in polycarbonate cages (47.6 × 26.0 × 20.3 cm, RC88D-UD, Alternate Design Mfg and Supply, Siloam Springs, Arizona, USA) with a bedding of wood shavings (Aspen chip, NEPCO, Warrensburd, NY, USA) and enrichments of a PVC tube, sizzle paper and nestlets. Rats were provided food (Prolab 2500 Rodent 5p14, Laboratory Animal Diet, LabDiet, St Louis, MO, USA) and tap water *ad libitum*. The housing environment consisted of a 12-hour light-dark cycle (light on at 7 am) and temperature and humidity of 23°C and 22%, respectively.

Animals were block randomised (list randomizer, random.org) by sex and strain to receive one of four treatments: 1) 800 mg/kg pentobarbital (PB, Euthanyl, 240 mg/mL, Bimeda-MTC Animal Health Inc., Cambridge, ON, Canada, pH of 11.018 ± 0.009 upon testing); 2) 800 mg/kg PB with 4 mg/kg lidocaine (PB+lido, Lidocaine Neat, 20 mg/mL, Pfizer Animal Health, Pfizer Canada Inc., Kirkland, QC, Canada); 3) Saline controls at 3.33 mL/kg (Sodium Chloride 0.9% Injection, FK Std., Fresenius Kabi Canada, Mississauga, ON, Canada; volume equal to PB) and 4) vehicle controls (SD: vehicle control pH of 11.0 at 3.33 mL/kg. Wistars: vehicle control pH of 12.5 at 3.33 mL/kg). [7] Vehicle controls were prepared as: propylene glycol (40%), ethanol (10%), water for injection and pH balanced to pH 11 or 12.5 (Chief Pharmacy, Calgary, AB, CAN and Chiron Compounding Pharmacy Inc., Guelph, ONT, CAN). Each injection was prepared in a 3 mL syringe with a 25 gauge 5/8” needle and 0.01 mL of blue food colouring added (Blue Food Colour, Club House, McCormick Canada, London, Canada). Rats were excluded and replaced if misinjection was confirmed at necropsy. Both experimenters (JR, CS) were blinded to the treatments and all assessments were performed between 7 am and 6 pm.

### Video recording (for behavioral assessments)

Three days before the experimental day, all rats were habituated daily to handling by the experimenters (JR and CS) and placement in the observation chamber (14 × 27 × 21 cm) for 10 minutes. During handling, both experimenters habituated the rat to the two-person injection technique: one experimenter (CS) cradled the rat in a backpack hold in dorsal recumbency with a 30° head down angle and extended the left pelvic limb. The other experimenter extended the right pelvic limb while simultaneously holding a capped hypodermic needle against the abdominal wall of the right caudal quadrant at a 45° angle to the body wall, as previously described. [6]

On the testing day, animals were weighed before placement into the observation box for baseline recording (Panasonic HC-V720P/PC, Panasonic Canada Inc., Mississauga, ON, Canada) for 10 minutes. Rats were then removed from the box and given a single intraperitoneal (IP) injection using the two-person injection technique. After injection (INJ), rats were immediately returned to the observation chamber for observation and video-recording until loss of righting reflex (LORR) occurred or until 10 minutes elapsed (whichever came first). At the first signs of ataxia, LORR was assessed by attempting to place the rat in left lateral recumbency, followed by dorsal recumbency. If the rat remained on its back for 10 seconds LORR was considered to be achieved. If the rat righted itself, LORR was reassessed every 30 seconds until achieved or up to 10 minutes. Following LORR, the animal was monitored for cessation of breathing (CB, visual assessment). At CB, the animal was placed in left lateral recumbency and monitored for cessation of heartbeat (CHB, thoracic auscultation with stethoscope). If CHB did not occur within 20 minutes of injection, the rat was euthanized with an overdose of inhaled isoflurane (IsoFlo®, Abbot Animal Health, Abbott Laboratories, North Chicago, IL, USA). Times to achieve LORR, CB and CHB were recorded. These physiologic data were collected for PB and PB+lido group animals.

### Behavioral assessments

Image collection for RGS scoring and behavioral assessments were performed by observers (JR and VL) blinded to treatment. For RGS scoring, an image was selected at three-minute intervals when the rats were not performing behaviors that could influence facial expressions (i.e. sleeping, sniffing, eating and grooming). Three images were collected during the 10 minute baseline video for each animal. For post-injection (PI) videos of animals that had LORR, three images were selected before LORR occurred (duration ranged from 56 to 105s). For PI videos in which LORR did not occur, three images were selected from each of the following intervals: first 80s of the video (average time to achieve LORR), from a 30s segment of the video (121 to 151s after IP injection, based on data showing LORR may not be achieved for up to 151s) [6] and during the last minute of observation (9 to 10 minutes after IP injection). Collected images were inserted into commercial presentation software (Microsoft PowerPoint, version 15.0, Microsoft Corporation, Redmond, WA, USA) and randomized with a macro (http://www.tushar-mehta.com/powerpoint/randomslideshow/index.htm). Each image was assessed as previously described. [11] Briefly, four action units (orbital tightening, ear changes, nose/cheek flattening and whisker changes) were scored from 0 to 2 (increasing score represents increasing pain). The following behaviors were assessed as relative frequencies: writhing and back arching. [12] These behaviors were identified over the first 151s of both BL and PI videos or before LORR. Writhing was defined as the contraction of the lateral abdominal walls where the abdomen appears concave. Writhing was also assessed before LORR or during the first 80s PI and during the last minute (9-10 minutes PI, where LORR did not occur). Back arching was defined as the arching of the back (with the abdomen pushed towards the ground or a vertical upwards arch).

### Necropsy

Skin was incised along the midline from the sternum to pubis and reflected back using blunt dissection. The linea alba was incised and the muscles along the costal arch were cut to expose the peritoneal cavity, which was photographed with viscera in place. The gastrointestinal tract from cardia to descending colon was then removed and any sections with blue staining opened to determine if staining was serosal or intraluminal. The liver, abdominal wall surrounding the injection site, and excised gastrointestinal tract were placed in 10% neutral buffered formalin solution for fixation of at least 7 days.

### Histological analysis

For histologic analysis, formalin-fixed samples of liver, gastrointestinal tract and abdominal wall muscle were embedded in paraffin, sectioned at 4 micrometer thickness and stained routinely with haematoxylin and eosin (H&E) stain. Samples were not collected from animals in which a misinjection was strongly suspected based on initial gross evaluation of abdominal contents. Slides were evaluated by a US board-certified veterinary pathologist (CGK) blinded to treatment for evidence of mesothelial (peritoneal) and submesothelial damage or inflammation.

### Statistical Analysis

Data were analysed with commercial statistical software (Prism 6.07, GraphPad Software, La Jolla, CA, USA). Normality was assessed with the D’Agostino-Pearson omnibus normality test. Physiologic data and RGS scores approximated a normal distribution while the relative frequency of writhing and back arching did not. An unpaired t-test was used to assess the differences between PB and PB+lido animals to achieve LORR, CB and CHB. Differences from baseline were assessed with a paired t-test, Wilcoxon test (for PB and PB+lido data), one-way ANOVA or a Friedman’s test (post-hoc Dunn’s, for saline and vehicle control data). Strains were analysed separately because of the different pH levels of the vehicle controls. Sample sizes were estimated (G*Power 3.1.9.2, Germany) for the two main behavioral outcomes: RGS and writhing. For the RGS, a sample size of 12 animals per group was estimated based on: alpha = 0.05, power = 0.8, SD = 0.25 and expected mean difference of 0.3. [11] For writhing, a sample size of 12 animals per group was estimated based on: alpha = 0.05, power = 0.8, SD = 4.74 and expected difference of 6. [12] A p-value of < 0.05 was considered statistically significant for all comparisons and 95% confidence intervals for the mean/median difference presented where available. Data in the figures are presented as mean ± SEM (RGS) or as median ± IQR (relative frequency of back arch and writhing). Data in text are presented as mean ± standard deviation. Data supporting the results are available in an electronic repository (Harvard Dataverse): XXX.

## Results

The misinjection rate was 25.2% (38/151), similar to previous studies. [6, 13–15] These rats were excluded from analysis. An additional rat was excluded due to a dosing error. The total number of SD females, SD males and Wistar females were n = 40, n = 36 and n = 36 respectively. As block randomization was maintained, these animals were divided equally into the four treatment groups: SD females (n = 10) SD males (n = 9) and Wistar females (n = 9). A maximum of 984 images could be captured for RGS assessment. The successful image capture rate was 99.7% with only three images that could not be captured from two rats because they were grooming during the majority of the observation period. Tissue samples were collected from 100 rats. Due to a planning error, formalin was unavailable for sample storage in 30 rats. Samples were not taken from 20 rats as misinjection was identified at initial necropsy. Samples from 17 rats were excluded because of misinjections identified during histological examination. There were 26 samples from SD male (PB and PB+lido: n = 4, saline: n = 7, VC: n = 11), 33 from SD female (PB, PB+lido and vehicle: n = 8, saline: n = 9) and 24 from Wistar female rats (PB and PB+lido: n = 3, saline and vehicle: n = 9).

There were increases in some behaviors, such as the RGS and writhing. These changes were only observed in animals that received vehicle controls and during the 151s and 10 mins PI periods.

### RGS

Increases in RGS scores were only observed in the rats that received the vehicle controls (SD: F (2.3, 41) = 8.8, p = 0.0004; Wistar: F (2.0, 16) = 6.4, p = 0.009; Fig. 1a, b). There were increases in RGS scores from BL after 10 min PI (SD: p = 0.007, 95% CI −0.77 to −0.12; Wistar: p = 0.024, 95% CI −0.91 to −0.074). Increases from BL at 151s PI was also observed in the SD rats (p = 0.0008, 95% CI −0.73 to −0.20) but not in the Wistar rats (p = 0.188, 95% CI −0.85 to 0.16). The mean of these scores also crossed a previously established intervention threshold of 0.67. [16] The RGS scores at 80s remained similar to BL (SD: p = 0.247, 95% CI −0.36 to 0.073; Wistar: p > 0.999, 95% CI −0.23 to 0.23).

**Fig 1.**
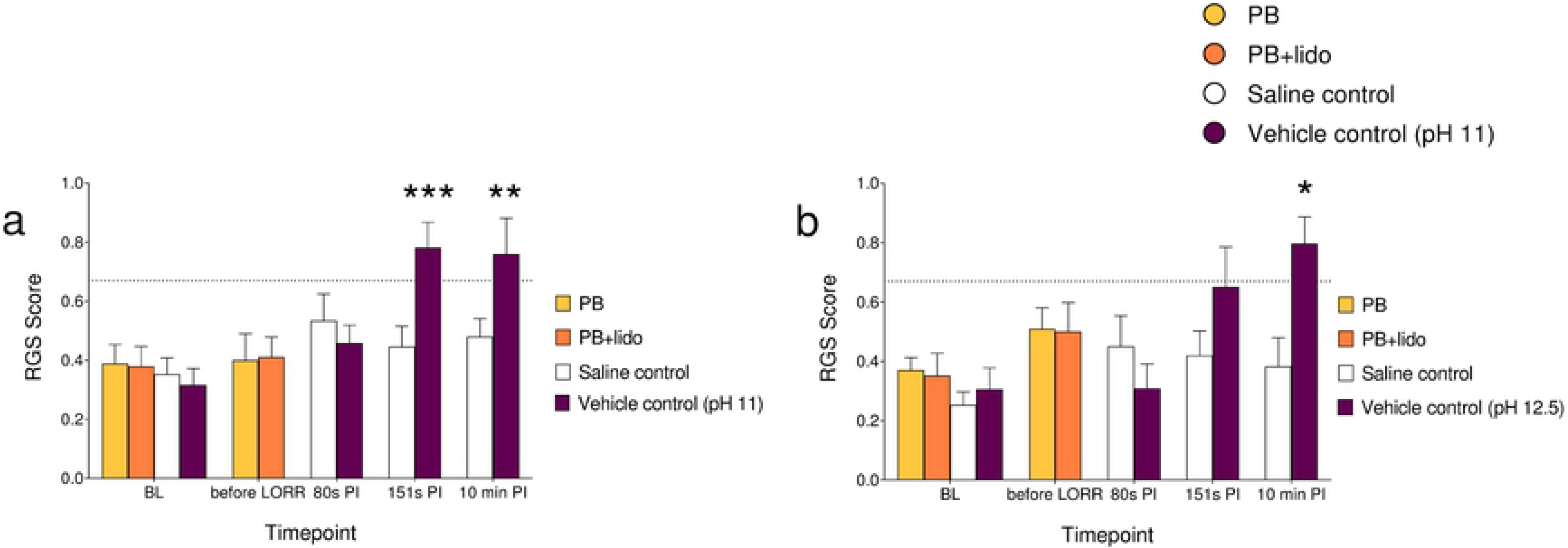
The RGS scores of rats that received sodium pentobarbital (PB), PB with lidocaine (PB+lido), saline controls (Control (SAL)) or vehicle controls (Control (pH 11 or 12.5)). (a) In the female and male Sprague Dawley group, significant increases from baseline were observed from the vehicle control group at the 151s and 10 min post-injection (PI) timepoints (p < 0.01). (b) In the female Wistar group, a significant increase from baseline was only observed at 10 min PI timepoint in the vehicle control group (p < 0.05). The horizontal dotted line represents a previously established threshold of 0.67. [16] Data presented as mean ± SEM. *p < 0.05, **p < 0.01, ***p < 0.001.

The PI RGS scores of animals which received PB (SD: p = 0.876, 95% CI −0.14 to 0.17; Wistar: p = 0.100, 95% CI −0.034 to 0.31), PB+lido (SD: p = 0.743, 95% CI −0.17 to 0.24; Wistar: p = 0.165, 95% CI −0.08 to 0.37) or saline (SD: F (2.4, 44) = 1.8, p = 0.173; Wistar: F (2.2, 18) = 1.3, p = 0.300) displayed similar scores to BL.

### Writhing

Similar to RGS scores, increases in the relative frequency of writhing behaviors were observed in the animals that received the vehicle controls (SD and Wistar: p < 0.0001; Fig. 2a, b). A higher relative frequency of writhing was observed at 151s PI compared to BL (SD: p < 0.00001; Wistar; p = 0.0004). However, at 80s (SD: p = 0.114; Wistar: p = 0.085) and 10 min PI (SD and Wistar: p > 0.999) the relative frequency of writhing was similar to BL. In PB, PB+lido and saline treatment groups, the relative frequency of writhing did not increase PI (PB; SD: p > 0.999; Wistar: p > 0.999. PB+lido; SD: p = 0.250; Wistar: p > 0.999. Saline controls; SD: p = 0.392; Wistar: p = 0.706, Fig 2a, b).

**Fig 2.**
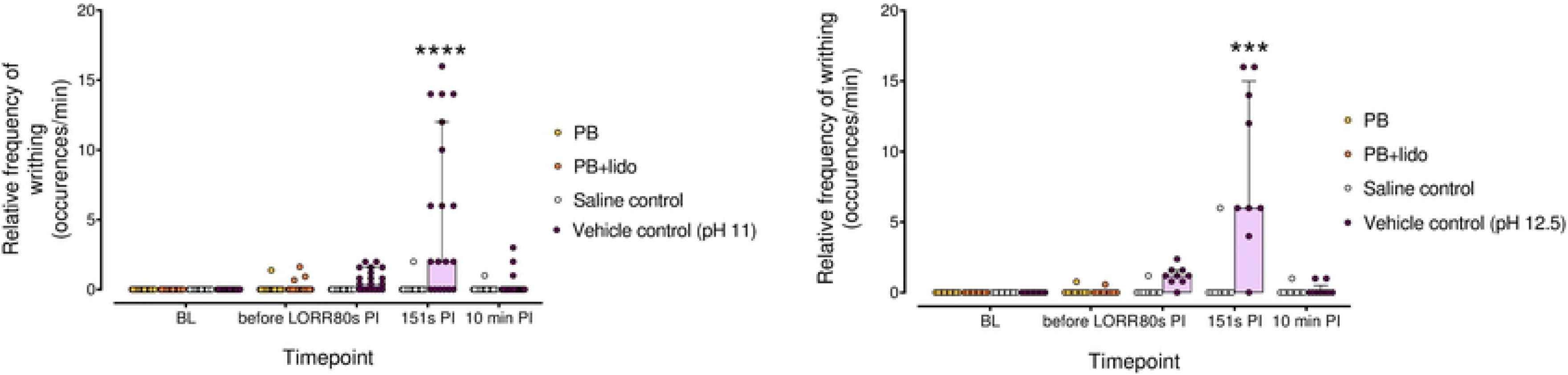
The relative frequency of writhing displayed by rats treated with sodium pentobarbital (PB), PB with lidocaine (PB+lido), saline controls (Control (SAL)) or vehicle controls (Control (pH 11 or 12.5)). Significant increases from baseline (BL) were only observed at the 151s post-injection (PI) timepoint in both female and male Sprague Dawley rats (p < 0.0001, a) and female Wistar rats (p < 0.001, b). Data presented as median ± IQR. ***p < 0.001, ****p < 0.0001.

Upon visual inspection of the data, a sex effect was apparent: SD males in the vehicle control groups displayed a lower relative frequency of writhing behavior in comparison to the female SD rats (Suppl. Fig. 1a, b). A higher relative frequency of writhing was observed in female SD rats that received vehicle controls at 151s PI only (p < 0.0001) but not in male rats (p = 0.249). The relative frequency of writhing was similar to baseline at all other timepoints in both SD males and females that received the vehicle controls (at 80s PI; males: p > 0.999, females: p = 0.113. at 10 min PI; males: p > 0.999, females: p = 0.896). Of animals that received saline, there were no differences from baseline in the SD males (80s, 151s and 10 min PI: p > 0.999). There were no significant differences from BL in writhing behavior in the PB and PB+lido groups (PB; female: p > 0.999; male: p = 1.00. PB+lido: female: p = 0.500; male > 0.999).

### Back arching

SD rats in the PB+lido and vehicle control groups expressed back arching more frequently during the PI period than at BL (PB+lido and vehicle control: p = 0.031; Fig. 3a, b). This increased frequency during PI was not observed in Wistar rats (PB+lido: p = 0.250, vehicle control: p = 0.063). For both SD and Wistar rats that received PB or saline, the relative frequency of back arching was similar to baseline (PB: SD; p = 0.250, Wistar; p > 0.999. Saline: SD; p = 0.313, Wistar: p > 0.999).

**Fig 3.**
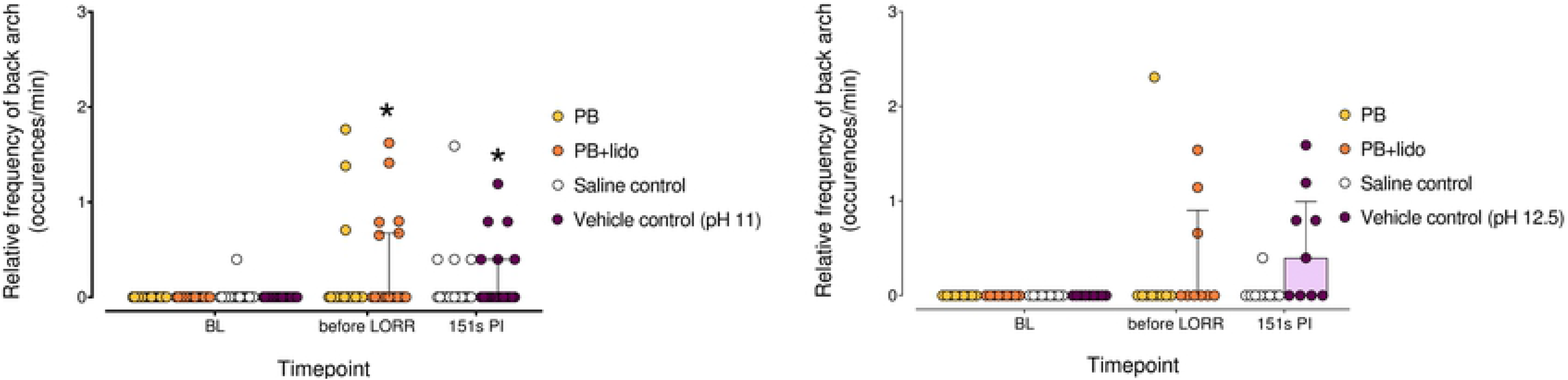
The relative frequency of back arching displayed by rats treated with sodium pentobarbital (PB), PB with lidocaine (PB+lido), saline control (Control (SAL)) or vehicle controls (Control (pH 11)). (a) With the female and male Sprague Dawley rats, significant increases from baseline (BL) were only observed in the PB+lido group before loss of righting reflex (LORR) and in the vehicle control group at the 151s PI timepoint (p < 0.05). (b) With female Wistar rats, there were no significant differences from BL at all timepoints and in the different treatment groups (p > 0.05). Data presented as median ± IQR. *p < 0.05.

Visual inspection of the data for sex differences was also performed and no differences were apparent (Suppl. Fig. 2a, b). Comparisons to BL did not reveal significant increases when the sexes were separated (PB; females: p = 0.500, males: p = 0.999. PB+lido; females: p = 0.125, males: p = 0.500. saline; females: p > 0.999, males p = 0.500. vehicle control; females: p = 0.125, males: p = 0.500).

### Physiologic data

There were no significant differences between animals in the PB and PB+lido treatment groups for each endpoint: INJ to LORR; PB: 78 ± 7.9s, PB+lido: 78 ± 11s, p = 0.91, 95% CI −4.71 to 5.28. LORR to CB; PB: 90 ± 19s, PB+lido: 94 ± 15s, p = 0.35, 95% CI −4.85 to 13.64. CB to CHB; PB: 84 ± 16s, PB+lido: 85 ± 17s, p = 0.73, 95% CI −7.30 to 10.30. INJ to CHB; PB: 252 ± 24s, PB+lido: 255 ± 20s, p = 0.64, 95% CI −9.13 to 14.7). Overall, the time between INJ to LORR, LORR to CB, CB to CHB and from INJ to CHB were 78 ± 9.2s, 92 ± 17s, 85 ± 16s and 254 ± 22s, respectively, for pooled data from PB and PB+lido groups. Twenty-five rats that received a misinjection of either PB or PB+lido achieved LORR at 149.9 ± 79.4s (range 74 to 347s).

### Histologic analysis

Each of the 83 slides included up to 7 representative sections of small and large intestine (3 gut sections: n = 9; 4 gut sections: n = 26; 5 gut sections: n = 33; 6 gut sections: n = 14; 7 gut sections: n = 1). The majority of slides included a section of liver (n = 81). Thirty-five included a section of abdominal wall and 12 included a section of pancreas attached to the associated duodenal segment. No evidence of mesothelial or submesothelial damage or inflammation was seen in any section aside from rare foci of mechanical trauma caused by the injection needle.

## Discussion

The results of this study show that: 1) IP injection of PB is painful and the source of pain is the alkali pH and 2) behaviors associated with pain are masked by the presence of PB.

As outlined in the CCAC [1] and AVMA [3] euthanasia guidelines, during the euthanasia of animals distress and pain must be minimized. The use of barbiturates, such as sodium pentobarbital, is designated as an acceptable method and preferred over other methods, such as inhalant anesthetics, because they are fast acting, inexpensive, readily available, have a long shelf life and are supposedly less aversive. [3] However, the methods to assess pain associated with IP PB have not been well defined. [3]

The highly alkali pH of PB solution (typically pH 11-12) has been suggested as a cause of pain when delivered IP. [1, 7] A few studies have reported that pain is present during IP PB injection because of changes in behaviors (i.e. increase writhing and reduction of locomotion and rearing, directed grooming [7, 8, 10, 17] levels of molecular markers (i.e. increase of spinal c-fos [9]) and the appearance of redness in the peritoneal cavity, indicative of inflammation. [10] Studies have also reported that writhing and spinal c-fos levels decreased when a local anesthetic, such as lidocaine or bupivacaine, was administered. [8–10] These results suggest that pain may be associated with IP PB injections. Unfortunately, not all of these studies have undergone peer review. [7, 10] Interpretation of changes in molecular markers alone is challenging as expression is altered by neuronal activation that may not be specific to nociception and nociception is not necessarily indicative of pain. [9, 18] Furthermore, short periods of nociceptive input, as seen during successful IP PB injection, are difficult to identify using changes in expression of many molecular markers. [18] Additionally, a failure to document a behavior in the presence of a drug causing sedation and reduced motor function, such as PB, should be interpreted cautiously. This could explain the apparent failure of the RGS to change following IP PB in one study. [8, 17] A novel approach, evaluating behavior directed at an alternative injection site (intra-plantar) reported low instances of paw licking, which were similar between mice injected with PB or saline. [17] Previous work has shown that loss of consciousness (as assessed with LORR) takes approximately 151 seconds to occur following injection (PB dose of 800 mg/kg) in rats. [6] This highlights the short window of opportunity for observations. This study was designed to clarify the role of pH in eliciting pain and explore the role of different behavioral outcomes in identifying pain over a relevant time frame.

Several behaviors have been used to study abdominal pain (often following laparotomy). [11, 12, 19] Of these, the grimace scales and writhing behavior were selected as they have also been shown to increase with exposure to a noxious substance injected IP and decrease with analgesics. [19, 20]. Back arching was reported to increase after laparotomy [12] but has not been specifically reported to increase in response to the IP injection of noxious substances in rodents. Unfortunately, these behaviors did not change reliably in this study (i.e. changes observed in PB or pH control groups).

### Writhing behavior

An increase in writhing behavior was observed only in the vehicle control groups at 151s PI and not in the PB groups. This differs from previous studies that reported an increase in writhing duration after IP injection of PB and the presence, though at a low incidence, of writhing after IP PB. [6, 8] These differences may result from the different methods of assessing writhing (relative frequency vs duration vs presence/absence). While writhing duration would be expected to be affected by the PB dose used (and consequent time to onset of sedation), the dose range employed by Khoo et al. [8] was similar (approximately 590-930 mg/kg) to that used in this study. Unfortunately, differences in methodology (duration versus relative frequency of writhing) preclude direct comparisons between these studies. We elected to use relative frequency to account for the different observation periods between individuals and treatment groups. Zatroch et al. [6] reported writhing in fewer than 50% of animals injected with either 200 or 800 mg/kg PB and observed writhing in a small number of animals (n = 2/9) receiving a saline control injection, highlighting the importance of this control. Interestingly, in the study reported here, writhing behavior was not sustained at the 10 minute observation period. This could reflect a reduction in pain over time or perhaps an increase in pain to a level that inhibited further writhing.

An unexpected effect of sex was observed, with male SD rats displaying fewer bouts of writhing than females. The source of this difference is unknown. However, this effect of sex was not maintained with the RGS scores, suggesting that other factors may have been involved.

### RGS

Similar to writhing behaviors, RGS scores only increased significantly in the vehicle control groups. In contrast to the writhing data, the RGS scores were increased at both 151s and 10 minute time points, and the average scores were close to or exceeded a previously established threshold associated with pain. [16] The maintenance of low RGS scores in the PB treated animals and increase in scores in the vehicle control group highlights the potential for agents with sedative/anesthetic properties to mask behavioral expression. This is a likely explanation for the failure of RGS scores to change in the study of Khoo et al. [8]

The results from both writhing and RGS observations indicate that an IP injection of PB is painful due to the alkali pH of the injectate. The low relative frequency of writhing and the low RGS scores in the PB groups support our hypothesis that the sedative effect of PB inhibits behavioral expression. Importantly, neither writhing nor RGS scores were increased at the 80s time point, suggesting that the onset of pain occurs after this time. Therefore, if loss of consciousness occurs rapidly, it is possible that pain may not be experienced following IP PB. This has important implications for situations in which time to unconsciousness is delayed, such as with misinjection and perhaps with lower doses of PB. [6] Animals that received a low dose and low volume of PB (200 mg/kg) took on average 25% longer to achieve LORR. [6] Our study also reported that rats which received a misinjection of PB take longer to achieve LORR. Unfortunately, it appears that misinjections are always possible, in part due to the variable location of the cecum. [6, 14, 15, 21]

### Histology

Histologic examination of the serosal surfaces of abdominal organs harvested from rats euthanized by IP injection does not offer information about pain associated with the procedure. Organs were evaluated for evidence of acute inflammation, including serosal and subserosal vasodilation and vascular congestion, neutrophilic margination and transmigration, mesothelial cell swelling, or mesothelial cell necrosis and sloughing. None of these features were seen, and no treatment group differences were detected by a blinded pathologist. This was not unexpected, as most histologic changes associated with acute inflammation take longer than 20 minutes to develop, which was the maximum interval between injection and death in this trial. The exception to this is vasodilation / active hyperemia, which can occur seconds to minutes after injury. Grossly visible peritoneal and serosal reddening has been previously described after IP PB injection, [10, 17] but this was not seen in this trial, however. Histologically, it is difficult to attribute significance to the presence or absence of red blood cells in blood vessels for several reasons. While the heart is still active blood may redistribute around the time of death through dysregulated autonomic control of vessels. After death, blood may redistribute under the influence of gravity. Blood can also be lost from vessels during tissue trimming and histologic processing. Finally, physiologic hyperemia due to digestion and peristaltic activity may cause intestinal sections to appear congested; this should not be interpreted as peracute inflammation. Therefore, although gross reddening of serosal surfaces caused by IP PB injection cannot be ruled out, it was not seen in this trial and was not supported histologically.

#### Limitations and future studies

A limitation of this study was the omission of a treatment group of a vehicle control combined with a local anesthetic. This is a necessary step to confirm if the use of a local anesthetic is effective in providing analgesia to counteract the pain associated with PB. This study only used one dose of PB. This dose was selected from previous work demonstrating benefits in terms of a faster death and reduced variability of effects. [6, 14, 15, 21]

In conclusion, IP injection of PB is painful, as indicated by the presence of behaviors associated with pain observed during vehicle control injections. Furthermore, these results highlight the importance of a vehicle control group and the limitations of interpreting behaviors in the presence of an agent with sedative properties, such as PB.

